# Structure of SARS-CoV-2 main protease in the apo state reveals the inactive conformation

**DOI:** 10.1101/2020.05.12.092171

**Authors:** Xuelan Zhou, Fangling Zhong, Cheng Lin, Xiaohui Hu, Yan Zhang, Bing Xiong, Xiushan Yin, Jinheng Fu, Wei He, Jingjing Duan, Yang Fu, Huan Zhou, Qisheng Wang, Jian Li, Jin Zhang

## Abstract

M^pro^ is of considerable interest as a drug target in the treatment of COVID-19 since the proteolytic activity of this viral protease is essential for viral replication. Here we report the first insight of the structure M^pro^ for SARS-CoV-2 in the inactive conformation under conditions close to the physiological state (pH 7.5) to an overall resolution of 1.9 Å. The comparisons of M^pro^ in different states reveal that substrate binding site and the active site are more flexible in the inactive conformation than that in the active conformations. Notably, compared with the active conformation of the apo state structure in pH7.6 of SARS, the SARS-CoV-2 apo state is in the inactive conformation under condition close to physiological state (pH7.5). Two water molecules are present in the oxyanion hole in our apo state structure, whereas in the ligand-bound structure, water molecular is absence in the same region. This structure provides novel and important insights that have broad implications for understanding the structural basis underlying enzyme activity, and can facilitate rational, structure-based, approaches for the design of specific SARS-CoV-2 ligands as new therapeutic agents.

## Introduction

Coronavirus Disease 2019 (COVID-19) caused by SARS-CoV-2 has been a global pandemic that severely threats to the global health and economy. However, there is currently no clinically approved vaccines or drugs against COVID-19 [1]. SARS-CoV-2 particles contain single, positive stranded RNA genome with a length of about 30 kb, with a 5’ cap structure and a 3’ poly (a) bundle. The SARS-CoV-2 genome encodes for replicase, spike glycoprotein (S), envelope protein (E), membrane protein (M) and nucleocapsid protein (N). SARS-CoV-2 main protease (M^pro^, also called 3C-like protease, 3CL^pro^) mediates the proteolytic processing of large replicase polyprotein 1a (pp1a) and pp1ab into non-structural proteins (NSPs) at 11 conservative sites. [2]. Thus, M^pro^ is of considerable interest as a drug target in the treatment of COVID-19 since the proteolytic activity of this viral protease is essential for viral replication. Mutational and structural studies have identified substrate binding site and active site of M^pro^ that confers specificity for the Gln-P1 substrate residue in the active conformation [3–5]. Structures of M^pro^ for SARS-CoV-2 has been solved in complexes with Chinese herbal and novel inhibitors very recently [6,7]. However, a structural description of these sites in the inactive conformation has remained elusive.

## Results

### Overall structure of M^pro^ for SARS-CoV-2

Here we report the first insight of the structure M^pro^ for SARS-CoV-2 in the inactive conformation under conditions close to the physiological state (pH 7.5) to an overall resolution of 1.9 Å (Table 1), guiding specific drug discovery and functional studies. The M^pro^ forms a dimer in the crystal and has two distinct dimer interfaces, which are located in the N-terminal domain (residues 1-11) and the oxyanion loop (residues 137-145) (Fig 1a). Comparison of our M^pro^ structure in the apo state to the previously reported M^pro^ structure in complex with inhibitor revealed a backbone (Cα) RMSD of 0.92 Å showing a similar overall structure [5–7] (Fig. 2a). As in ligand-bound M^pro^ structures, the protein consists of N-finger and other three domains that bind inhibitor at the cleft between domains I and II. N-finger (residues 1-7) is a loop located in the dimer interface and involved in the N-terminal auto cleavage. The domain I (residues 8-101) is comprised of three small α-helices and six β-strands. The domain II (residues 102-184) consists of six β-strands. The domain III is composed of five β α-helices, which are closely related to the proteolytic activity.

**Table. 1.** Statistics for data processing and model refinement of COVID-19 M^pro^.

|  |  |
| --- | --- |
| PDB code | 7C2Q |
| Synchrotron | SSRF |
| Beam line | BL17U1 |
| Wavelength (Å) | 0.97918 |
| Space group | P2<sub>1</sub>2<sub>1</sub>2<sub>1</sub> |
| *a*, *b*, *c* (Å) | 67.95, 102.66, 103.54 |
| $\alpha$, $\beta$, $\gamma$ (°) | 90.00, 90.00, 90.00 |
| Total reflections | 716573 |
| Unique reflections | 55189 |
| Resolution (Å) | 1.93(1.98-1.93) |
| R-merge (%) | 11.8(149.3) |
| Mean *I* / $\sigma$ (*I*) | 15.4/2.2 |
| Completeness (%) | 99.9(99.9) |
| Redundancy | 13.0(12.8) |
| Resolution (Å) | 72.90-1.93 |
| *R*<sub>work</sub>/*R*<sub>free</sub>(%) | 22.63/26.61 |
| Atoms | 4917 |
| Mean temperature factor (Å<sup>2</sup>) | 36.7 |
| Bond lengths (Å) | 0.008 |
| Bond angles (°) | 0.88 |

260 Values in parentheses are for the highest-resolution shell.

**Fig. 1.**
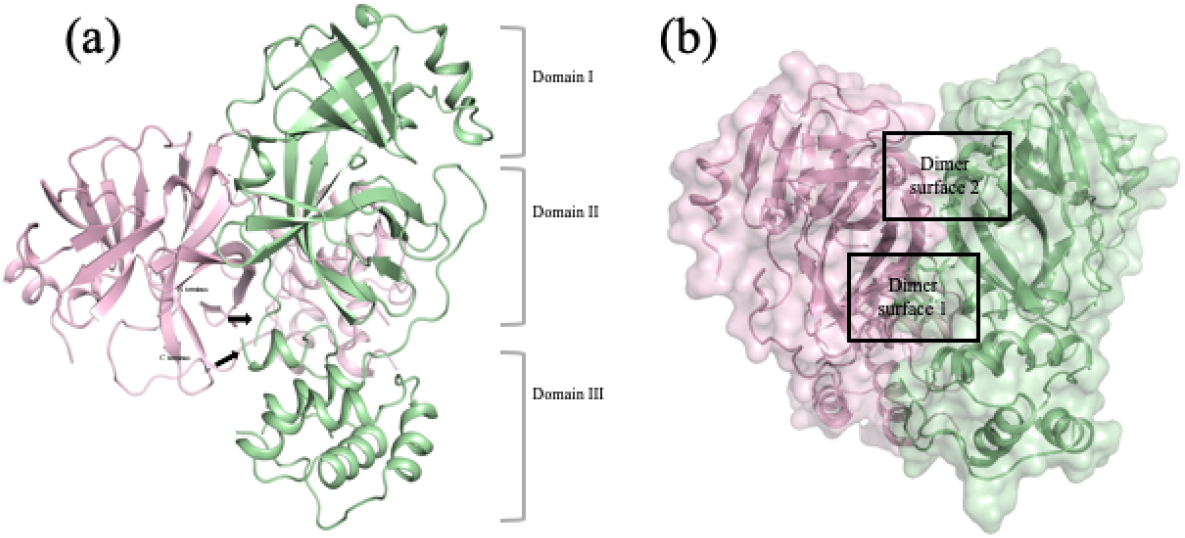
The apo state structure of M^pro^ of SARS-CoV-2 in the inactive conformation. (a) The structure of the M^pro^ dimer is shown in stereo. Individual protomers are shown in red and green. (b) Two dimer interfaces of M^pro^ of SARS-CoV-2. Dimer interface 1 and 2 are located in the oxyanion hole and N-terminal domain, respectively. (c) Comparison of M^pro^ structures in the apo state of (green) in SARS-CoV-2, with inhibitor N3 in SARS-CoV-2 (red, PDB: 6LU7), with inhibitor 11b in SARS-CoV-2 (orange, PDB: 6LZE) and in the apo state of SARS (gray, PDB:1UJ1). N3 and 11b are shown in pink and cyan, respectively. (d) Comparison of substrate binding site and active site in the apo state (green) and in the ligand-bound state in M^pro^ of SARS-CoV-2. 11b are shown in cyan. (e) Structure of M^pro^ bound with 11b in an active conformation. (f) Structure of M^pro^ in an inactive conformation. Water 1 and 2 are shown in red spheres.

**Fig. 2.**
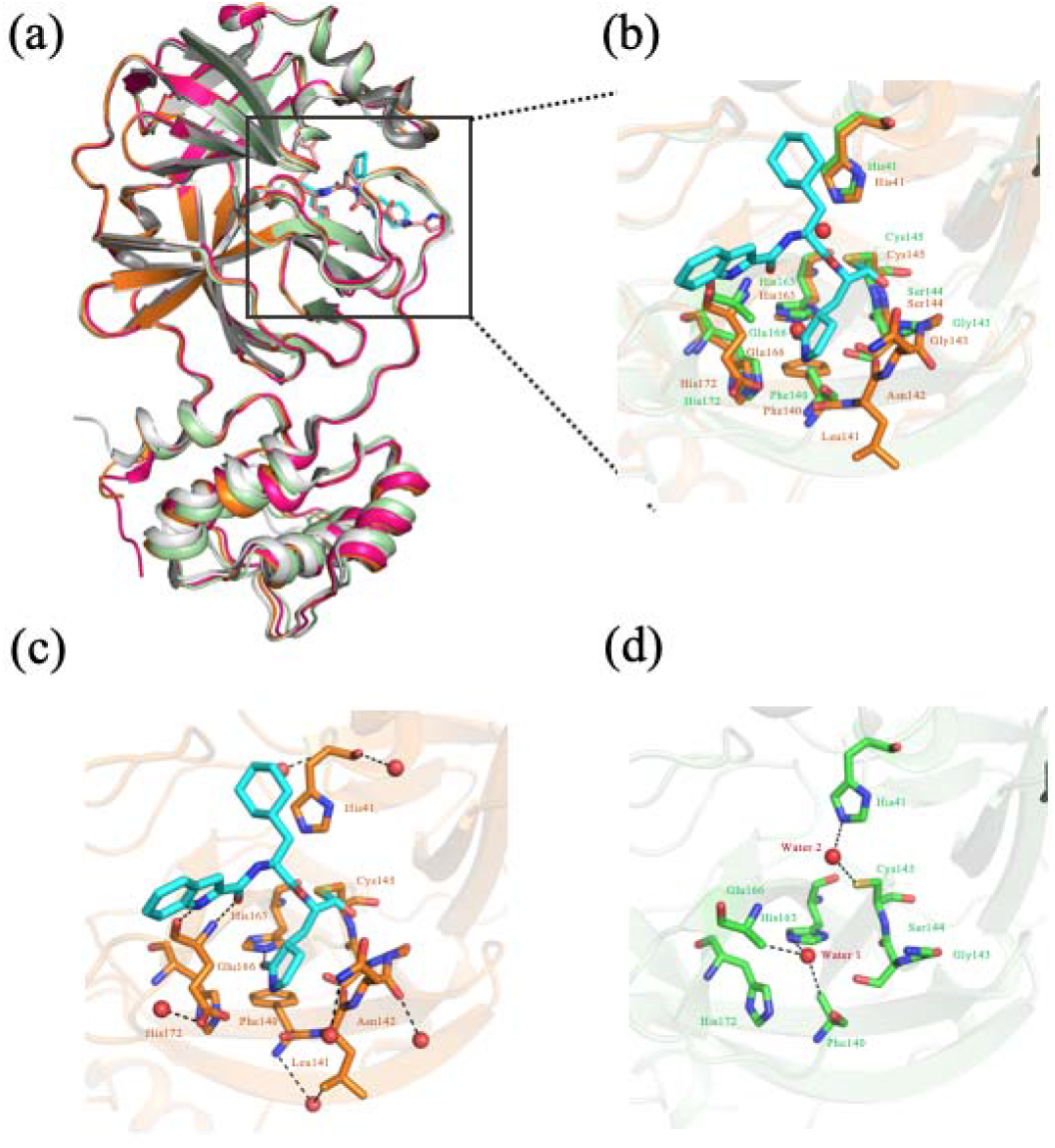
(a) Comparison of M^pro^ structures in the apo state of (green) in SARS-CoV-2, with inhibitor N3 in SARS-CoV-2 (red, PDB: 6LU7), with inhibitor 11b in SARS-CoV-2 (orange, PDB: 6LZE) and in the apo state of SARS (gray, PDB:1UJ1). N3 and 11b are shown in pink and cyan, respectively. (b) Comparison of substrate binding site and active site in the apo state (green) and in the ligand-bound state in M^pro^ of SARS-CoV-2. 11b are shown in cyan. (c) Structure of M^pro^ bound with 11b in an active conformation. (d) Structure of M^pro^ in an inactive conformation. Water 1 and 2 are shown in red spheres.

### Structural comparisons between M^pro^ in the apo states and other M^pro^ structures

There were, however, several notable local differences between the apo and ligand-bound structures. Electron density of the N-finger (residues 1-2), oxyanion loop (residues 141-142), C-terminal domain (resides 299-306) were insufficient for backbone tracing, suggesting the flexibility of this region in the apo state. In addition, electron densities of side chains Phe140 and Glu166, which are key residues involved in the substrate binding are missing at this high resolution that may reflect different conformation of the apo state (Fig. 2b).

The oxyanion hole composed of backbone amides or positively charged residues is directly related to the enzyme activity and substrate binding. In ligand-bound structures of M^pro^, the oxyanion hole consists of loop (residues 140-145), negatively charged residues Glu166, positively charged residues His41, His163 and His172 remains in an active conformation (Fig. 2c) [6]. A π-π stacking interaction (Phe140/His163) is found in the oxyanion hole. A hydrogen bond and salt bridge involving Glu166 with water and His172 at the domain II further stabilize oxyanion hole. However, the oxyanion loop (residues 137-145) is less well ordered and the side chains of Glu166 and Phe140 cannot be fit well due to poor density in our apo state structure. The salt bridge and π-π stacking interactions between Glu166/His172 and Phe140/His163 are broken, resulting in rearrangements in this region and further collapses of the oxyanion hole (Fig. 2d).

### M^pro^ in the apo states at pH 7.5 is in an inactive conformation

We propose that M^pro^ in the apo state is in the inactive conformation in the physiological condition, which is different from the active conformation of ligand-bound structures of M^pro^ [5]. The disordered N-finger is another feature of the inactive conformation. N-finger plays an important role in the formation of the active site and auto cleavage activity of M^pro^ [4]. Gly2 has interactions with Gly143 in the oxyanion loop in the neighboring protomer, stabilizing the active site and dimer in the active conformation, while the electron density of Gly2 is completely missing in the apo state. Inactive conformation in the apo state of our structure is further supported by the flexibility of N-finger in the apo state. It is consistent with the proof that lack of N-finger in TGEV M^pro^ is almost completely inactive [8]. Interestingly, His163 forms hydrogen bonds with water molecular (Water 1) in our structure, which is not observed in the active conformation. Another unprecedent water molecular (Water 2) is found at Cys145-His41 catalytic dyad in the active site, working as bridge for proton transfer. We speculate that these water molecules may affect negatively charged oxygen of the substrate or inhibitor, which suffers from steric hindrance, making rational drug design more difficult (Fig. 2b and d).

## Discussion

In summary, we determined the apo state structure of M^pro^ for SARS-CoV-2 in the inactive conformation. The comparisons of M^pro^ in different states reveal that substrate binding site and the active site are more flexible in the inactive conformation than that in the active conformations. Notably, compared with the active conformation of the apo state structure in pH7.6 of SARS, the SARS-CoV-2 apo state is in the inactive conformation under condition close to physiological state (pH7.5). The instable and disordered regions of oxyanion hole and the active site in the inactive conformation will raise the activation energy of the protease necessary for the reaction, slow down catalysis and finally extend the replication cycle of the virus. These structural differences may reveal the underlying reasons of why some patients infected with SARS-CoV-2 have longer virus latency than that of SARS. Further studies of detailed molecular mechanisms of SARS-CoV-2 pathogenesis are needed. For the drug design based on the structure, water molecules imbedded in the oxyanion hole and corresponding interactions should be taken into more consideration. Two water molecules are present in the oxyanion hole in our apo state structure, whereas in the ligand-bound structure, water molecular is absence in the same region. The water molecules, which is found near His163 and His41 in the occluded pocket, stabilizes the positively charged His residues, increasing the steric hindrance that may slow down the enzyme reaction and decrease the catalytic efficiency of the enzyme. Altogether, the apo state structure of M^pro^ for SARS-CoV-2 is an important complementary to the available structures. This structure provides novel and important insights that have broad implications for understanding the structural basis underlying enzyme activity, and can facilitate rational, structure-based, approaches for the design of specific SARS-CoV-2 ligands as new therapeutic agents.

## Materials and Methods

### Protein purification and crystallization

The cDNA of full length COVID-2019 main protease 3CL (NC_045512) were was optimized and synthesized (Generay, China) connected into vector pET28a to obtain the wanted plasmid. The plasmid was transformed into competent cell E.coli Rosetta DE3. The bacteria were grown in 800mL of LB (Luria-Bertani) broth at 37°C. When the OD600 reach 0.6-0.8, 500μM IPTG was added to induce the E. coli expression and then incubated 3-5h at 30°C. Centrifuge the cells at 10000g for 10min at 4 °C, discard the supernatant, collect the precipitate and add buffer A(100mM Tris/HCl buffer, pH7.5,300mM NaCl 10mM imidazole and 5% glycerol) to blend the collected cells, which were broken up by JNBIO 3000 plus(JNBI). The supernatant containing needed protein was acquired by centrifugation at 30000g, 4°C for 30min. Transfer the supernatant into 5ml Ni-NTA Ni2+-nitrilotriacetate column (GE healthcare) and the protein wanted was loaded onto the column. Add buffer B(100mM Tris/Hcl buffer, pH7.5, 300mM NaCl, 100mM imidazole, and 5% glycerol) into beads which use imidazole to wash. The His tagged protein was eluted by buffer C (50 mM Tris-HCl pH 7.5, 300 mM NaCl and 300 mM imidazole). Superdex 200 PG gelfiltration column (GE healthcare) can more purify protein and remove imidazole, while need to change the buffer to buffer C(25mM HEPES buffer, pH7.5, 300mM NaCl, 2mM DTT and 5% glycerol). Collect postive peak protein and use tiny part test by SDS-PAGE. In the end, the protein was flash-frozen in liquid nitrogen and stored at −80°C.

Thaw the protein and concentrate it to 5mg/ml in Amicon Ultra-15,10000Mr cut-off centrifugal concentrator (Millipore). The hanging drop vapor diffusion method was useful to gain crystal at 4°C. The crystals were grown with buffer containing 0.1M HEPES sodium 7.5, 10% Propanol and 20% PEG4000 in 3-5 days.

### Data collection, structure determination and refinement

The crystals were tailored with cryo-loop (Hampton research, America)and then flash-frozen in liquid nitrogen to collect better X-ray data. All data sets were collected at 100 K on macromolecular crystallography beamline17U1 (BL17U1) at Shanghai Synchrotron Radiation Facility (SSRF, Shanghai, China). All collected data were handled by the HKL 2000 software package. The structures of COVID-2019 main protease 3CL were determined by molecular replacement with PHENIX software. The of 3T0H was referred as a model. The program Coot was used to rebuild the initial model. The models were refined to resolution limit 1.93Å by using the PHENIX software. The superimposed data was analyzed with PyMOL software package. The complete wanted data collection and statistics of refinement are shown in Table 1. The structure has been deposited in PDB (PDB code 7C2Q).

## Acknowledgments

Jin Zhang was supported by the Thousand Young Talents Program of China, the National Natural Science Foundation of China (grant no. 31770795; grant no. 81974514), and the Jiangxi Province Natural Science Foundation (grant no. 20181ACB20014). Jian Li was supported by the Open Project of Key Laboratory of Prevention and treatment of cardiovascular and cerebrovascular diseases, Ministry of Education (No. XN201904), Gannan Medical University (QD201910) and Jiangxi“Double Thousand Plan”. This work was also supported by Ganzhou COVID-19 Emergency Research Project.

## Author contributions

Jin Zhang and Jian Li designed the project. Xuelan Zhou, Fanglin Zhong, Xiaohui Hu and Cheng Lin made constructs for expression and determined the conditions used to enhance protein stability. Huan Zhou and Qisheng Wang carried out X-ray experiments, including data acquisition and processing. Jian Li and Jin Zhang built the atomic model. Jin Zhang, Jian Li and Jingjing Duan drafted the manuscript. All authors contributed to structure analysis/interpretation and manuscript revision. Jin Zhang and Jian Li initiated the project, planned and analyzed the experiments, and supervised the research.

